# Resistance to amitraz in the parasitic honey bee mite *Varroa destructor* is associated with mutations in the β−adrenergic-like octopamine receptor

**DOI:** 10.1101/2021.08.06.455389

**Authors:** Carmen Sara Hernández-Rodríguez, Sara Moreno-Martí, Gabrielle Almecija, Krisztina Christmon, Josephine D. Johnson, Marie Ventelon, Dennis vanEngelsdorp, Steven C. Cook, Joel González-Cabrera

## Abstract

*Varroa destructor* is considered a major reason for high loss rate of Western honey bee (*Apis mellifera*) colonies. To prevent colony losses caused by *V. destructor* it is necessary to actively manage the mite population. Beekeepers, particularly commercial beekeepers, have few alternative treatments other than synthetic acaricides to control the parasite, resulting in intensive treatment regimens that led to the evolution of resistance in mite populations.

To investigate the mechanism of the resistance to amitraz detected in *V. destructor* mites from French and U.S. apiaries, we identified and characterized octopamine and tyramine receptors (the known targets of amitraz) in this species. The comparison of sequences obtained from mites collected from different apiaries with different treatment regimens, showed that the amino acid substitutions N87S or Y215H in the OctβR were associated with treatment failures reported in French or U.S. apiaries, respectively. Based on our findings, we have developed and tested two high throughput diagnostic assays based on TaqMan^®^ able to accurately detect mites carrying the mutations in this receptor. This valuable information may be of help for beekeepers when selecting the most suitable acaricide to manage *V. destructor*.

## INTRODUCTION

The ectoparasitic mite *Varroa destructor* (Anderson and Trueman), shifted hosts from the Eastern honeybee (*Apis cerana* L.) to the Western honey bee (*Apis mellifera* L.) in the late 1950’s (Traynor et al. 2020). Since then, it has spread almost exclusively as clonal lineages throughout the world (Solignac et al. 2005). In *A. cerana, V. destructor* causes little damage to the colonies since the parasite’s population growth is limited as mites can only reproduces in drone brood, which are only available in large numbers early in summer. In contrast, *V. destructor* successfully reproduces in both drone and worker brood of *A. mellifera* (Beaurepaire et al. 2015). *Varroa destructor* damages the host by feeding directly on the fat bodies, by vectoring viruses (Boecking and Genersch 2008; Ramsey et al. 2019) and reducing natural defences (Aronstein et al. 2012). If left unmanaged, *V. destructor* will kill the colonies within a few years (Martin et al. 1998). This mite is considered one of the major causes for seasonal colony losses of the Western honey bee (Steinhauer et al. 2018).

Beekeepers have an assortment of chemical and non-chemical methods to implement Integrated Pest Management (IPM) strategies for controlling *V. destructor*. Most of beekeepers use synthetic chemicals to treat their colonies, since they are easier to use and appear to be most effective and consistent at reducing losses (Rosenkranz et al. 2010; Haber et al. 2019). Globally, the most commonly registered acaricides are the pyrethroids flumethrin and tau-fluvalinate, the organophosphate coumaphos, and the formamidine amitraz. In the past, tau-fluvalinate and coumaphos have been the most widely used treatments, but now these pesticides are less effective. The intensive use of pyrethroids to control *V. destructor* since the 1980’s resulted in the independent emergence of resistance to these chemicals in mite populations from Europe and North America (Milani 1995; Elzen et al. 1998; Mozes-Koch et al. 2000; Sammataro et al. 2005; Gracia-Salinas et al. 2006; Kim et al. 2009; González-Cabrera et al. 2013; Hubert et al. 2014; González-Cabrera et al. 2016; González-Cabrera et al. 2018). Coumaphos was brought to market as an alternative varroacide treatment, but overuse of this product also resulted in the evolution of resistance (Elzen and Westervelt 2002; Maggi et al. 2009; Maggi et al. 2011). Moreover, residues of varroacides persist and accumulate in beeswax (Bonzini et al. 2011; Calatayud-Vernich et al. 2018; Traynor et al. 2020), posing a sublethal threat to honey bees (Desneux et al. 2007) and possibly maintaining the selection pressure on mite populations, so preventing resistance reversion, as already reported for pyrethroids resistance in *V. destructor* (Milani and Della Vedova 2002; Medici et al. 2016; González-Cabrera et al. 2018; Mitton et al. 2018).

To delay the evolution of resistance, rotation of products with different modes of action is recommended (IRAC; https://www.irac-online.org/), but the lack of effective alternatives makes chemical rotation a non-practical solution for beekeepers. As a result, beekeepers are over reliant on amitraz to control mites (Haber et al. 2019), which would select for resistant mites and it may explain consistent field reports of reduced miticidal (Elzen et al. 1999; Elzen et al. 2000; Rodríguez-Dehaibes et al. 2005; Maggi et al. 2010; Kamler et al. 2016; Rinkevich 2020).

In *V. destructor*, the mechanism of resistance to pyrethroids is already known. It is caused by substitution of key residues within the voltage gated sodium channel (VGSC), the major target site for pyrethroids (González-Cabrera et al. 2013; Hubert et al. 2014; González-Cabrera et al. 2016; González-Cabrera et al. 2018). Regarding the resistance to coumaphos, studies carried out with other species have reported that it may be associated with either mutations in its target site, the enzyme acetylcholinesterase, duplication of the acetylcholinesterase gene, or with alterations in the expression of detoxification enzymes (Feyereisen et al. 2015). However, in *V. destructor*, the mechanism(s) involved in the resistance to coumaphos remains unclear. The downregulation of a cytochrome P450 involved in the activation of coumaphos have been described as associated with the resistance reported in mites collected from the Greek island of Andros (Vlogiannitis et al. 2021).

In insects and acari, amitraz binds to the receptors of octopamine and tyramine (Kumar 2019). The octopamine (OAR) and the tyramine (TAR) receptors belong to the superfamily of G-protein coupled receptors (GPCRs). GPCRs are known to be involved in recognizing extracellular messengers, transducing signals to the cytosol, and mediating the cellular responses necessary for the normal physiological functions of organisms (Liu et al. 2021). Octopamine and tyramine receptors are classified as α-adrenergic-like octopamine receptors (Octα_1_Rs and Octα_2_Rs), β-adrenergic-like octopamine receptors (Octβ_1_Rs, Octβ_2_Rs, and Octβ_3_Rs), and tyramine receptors (TAR1, TAR2, TAR3) (Finetti et al. 2021).

Uncovering the molecular mechanisms involved in resistance to pesticides is essential for rapid detection and for designing effective management approaches. In this study, we identified and characterized octopamine and tyramine receptors of *V. destructor*. Two amino acid substitutions in Octβ_2_R associated with reported field treatment failures of amitraz in France and the U.S. were identified. Finally, two robust high throughput diagnostic assays were developed to identify *V. destructor* mites carrying these mutations in order to aid in resistance management in affected communities.

## MATERIAL AND METHODS

### *V. destructor* samples

Samples reporting failures after treatment with amitraz in France were collected in 2019 from five apiaries belonging to departments 38 (Isère), 42 (Loire), and 63 (Puy-de-Dôme). These apiaries have been treated with amitraz for several years in a row. Mites were collected from capped brood at the end of treatment with amitraz (70 days after the application of strips) and stored at -20 °C until used for molecular analysis. Mites collected at departments 4 (Alpes-de-Haute-Provence), 26 (Drôme), and 49 (Maine-et-Loire) were not treated with amitraz for at least one year before collection.

U.S. samples were collected as part of different surveys and research efforts not specifically designed for identification of the mechanism of resistance to amitraz (Table S1). Bee Informed Partnership Inc. (BIP) conducted a field trial in the fall of 2018 to test the efficacy of the product Apivar^®^ (a.i. amitraz) to reduce *V. destructor* mite infestation in colonies from active commercial beekeeping operations in the U.S. The trial was conducted within 2 commercial beekeeping operations from 2 different geographic regions. A total of 72 colonies (12 colonies per yard, in 3 yards for each operation) were followed over 42 days after treatment. In each yard, half of the colonies were treated with Apivar^®^ while the other half received a positive control product, Apilife Var^®^ (a.i. thymol). *Varroa destructor* load was estimated by a lab wash of a sample of ∼300 bees collected from a brood frame (Dietemann et al. 2013).

Phoretic mites from the BIP project were collected from colonies taking part in field trials conducted in Oregon and Michigan in 2018. The mites collected were those still in the colony while treatments were ongoing, and so survived at least a partial treatment exposure. Of mites from the 72-colony trial, we randomly chose mites from four colonies being treated with Apivar^®^ and four colonies treated with Apilife Var^®^ as positive control. We also analysed some mites collected in 2020 as part of the U.S. National Honey Bee Disease Survey (NHBDS). We looked at samples collected from Delaware, Massachusetts, Montana and Pennsylvania. Mite samples previously used to detect tau-fluvalinate resistance in U.S. mite populations and collected from these same states but from 2016 and 2017 NHBDS efforts, were also used (Millán-Leiva et al. 2021a). Samples from New Jersey were sent by a New Jersey state apiarist and were collected from apiaries reporting amitraz failure in 2018 (Styles, Personal communication).

Susceptible samples were collected from 2016 to 2019 in apiaries without exposure to amitraz from Iran, New Zealand, Spain, and the UK.

### Evaluation of amitraz efficacy

Acaricide efficacy of the amitraz treatments in MM16 and J11 French apiaries were calculated according to the Guideline on veterinary medicinal products controlling *Varroa destructor* parasitosis in bees (EMA 2010). Amitraz strips were introduced into hives at day 1 and they were removed at day 70. The number of mites in the inspection boards was registered every two days along the treatment. The residual number of mites was determined with a follow-up treatment using oxalic acid at day 91 and the final count of the dead mites at day 106. The treatment efficacy (E) was calculated as % of mite reduction as follows:

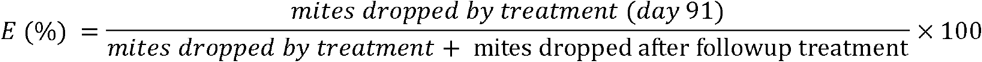

### Identification of receptors and phylogenetic analysis

Analysis of the contigs resulting from a transcriptomic analysis of *V. destructor* were previously carried out by our laboratory (BioProject ID PRJNA531374), allowing us to annotate putative octopamine-like receptors. The identity of these sequences was validated after searching in the *V. destructor* genome (BioProject PRJNA413423) (Techer et al. 2019) via BLASTn. Further comparison with previously annotated octopamine and tyramine receptors from related arthropod species was also carried out via multiple sequence alignment (47 sequences of octopamine and tyramine receptors were used, see Table S2). Protein alignments, tree generation for the phylogenetic analysis, electropherogram editions and sequence assembling were conducted using Geneious software (Geneious version 9.1.5 (http://www.geneious.com (Kearse et al. 2012)). Figures representing protein alignments were generated using CLC Sequencer Viewer 6.8.1. (www.clcbio.com).

### Amplification and sequencing of receptor cDNAs

Pools of 5 mites were ground to powder in liquid nitrogen and total RNA was extracted using the RNeasy Mini Kit (Qiagen) according to the manufacturer’s recommendations. RNA (0.5 – 1 µg) was reverse transcribed to cDNA using Maxima H minus First Strand cDNA synthesis kit (ThermoFisher Scientific) using oligo dT_18_ (250 ng). First strand cDNA was used as a template for PCR. Amplification of *Vd_octα*_*2*_*r* Open Reading Frame (ORF) was conducted using primers Vd_OctAR_5UTR and Vd_OctAR_3UTR. Amplification of *Vd_octβ*_*2*_*r* ORF was done using primers Vd_OctBR_5UTR1 and Vd_OctBR_3UTR. Amplification of *Vd_tar1* ORF was carried out using primers Vd_TAR1_5F and TAR1_3R (Table S3). For PCR amplifying the ORFs, 1 µl of cDNA was mixed with 100 ng of each primer, 25 µl of DreamTaq Green PCR Master Mix (ThermoFisher Scientific) and water to a final volume of 50 µl. Cycling conditions were: 94 °C for 2 min followed by 35 cycles of 94 °C for 45 s, 60 °C for 45 s and 72 °C for 2 min, and final extension at 72 °C for 5 min. The PCR fragments were purified using the NucleoSpin™ Gel and PCR Clean-up Kit (Thermo Scientific) and sequenced (Stabvida, Portugal) using the sets of primers showed in Table S3.

### Genomic DNA sequencing

DNA was extracted from individual mites using DNAzol^®^ reagent (ThermoFisher Scientific) following the manufacturer’s protocol. Primers used to PCR amplify and sequence the octopamine and tyramine receptor genes are described in Table S3. The mutation at position 260 of *octβ*_*2*_*r* was screened by amplifying the genomic region flanking the mutation site with primers Vd_OctBR_5UTR3 and Vd_OctBR_563R. The flanking region of the mutation at position 643 of *octβ*_*2*_*r* was amplified with primers Vd_OctBR_476F and Vd_OctBR_437iR. The PCR conditions were similar to those described above except for the extension step, which was run at 72 °C for 1 min. PCR amplicons were purified and sequenced as described above.

### Protein structure simulation

The online server for protein structure prediction I-TASSER (Yang and Zhang 2015) was used to generate a theoretical three-dimensional structure of *V. destructor* Octβ_2_R and TAR1. From the default settings of I-TASSER, the structure conformation with higher C-score for each receptor was chosen. C-score is typically in the range of [-5, 2], where a C-score of a higher value indicates a model with a higher confidence and vice-versa. The topology of the Octβ_2_R receptor in the membrane was represented using the webservice PROTTER (Omasits et al. 2014), which uses Phobius (Kall et al. 2004) for prediction of transmembrane topology and the N-terminal location. The predictions about the effects of the mutations in the receptor were obtained with SNAP2 (Hecht et al. 2015), PolyPhen2 (Adzhubei et al. 2010), I-Mutant2.0 (https://folding.biofold.org/i-mutant/i-mutant2.0.html), and HOPE (Venselaar et al. 2010).

### TaqMan^®^ diagnostic assays

The sequence of the *Vd_octβ*_*2*_*r* gene described in this study was used to design primers (flanking the N87 and Y215 positions in the Vd_Octβ_2_R protein) and two minor groove-binding probes (MGB) (ThermoFisher Scientific) using the Custom TaqMan^®^ Assay Design Tool (https://www.thermofisher.com/order/custom-genomic-products/tools/genotyping/). For the detection of N87S mutation, forward OctR_Vd_87_F (5’-CGCCCTGTTCGCGATGA-3’) and reverse OctR_Vd_87_R (5’-ATCCACTTGCCCGAAATGGT-3’) primers (standard oligonucleotides with no modification) were used. The probe Vd_N87S_V (5’-ACGACGCATTGAATG-3’) was labelled with the fluorescent dye VIC^®^ for the detection of the wild-type allele, and the probe Vd_N87S_M (5’-CGACGCACTGAATG-3’) was labelled with the fluorescent dye 6FAM™ for detection of the N87S mutation. For the detection of Y215H mutation, forward OctR_Vd_215_F (5’-GGATACCGTGCTCAGTAATGCT-3’) and reverse OctR_Vd_215_R (5’-CTGTCGGGTCGCTTCTAGATAG-3’) primers (standard oligonucleotides with no modification) were used. The probe Vd_Y215H_V (5’-ATGCGCCAATAAGTGAAT-3’) was labelled with the fluorescent dye VIC^®^ for the detection of the wild-type allele, and the probe Vd_Y215H_M (5’-CGCCAATGAGTGAAT-3’) was labelled with the fluorescent dye 6FAM™ for detection of the Y215H mutation. Each probe also had a 3’non-fluorescent quencher and a minor groove binder at the 3’ end. This minor groove binder increases the Tm between matched and mismatched probes providing more accurate allele discrimination (Afonina et al. 1997). Genomic DNA extraction from adult mites and TaqMan^®^ assays were carried out as described by González-Cabrera et al. (2013) using a StepOne Real-Time PCR System (ThermoFisher Scientific).

## RESULTS

### Identification of *V. destructor* octopamine and tyramine receptors

Manual curation of transcriptomic data obtained in our laboratory (BioProject ID PRJNA531374) showed that a few contigs contained sequences likely belonging to G protein-coupled receptors (GPCR) and more specifically to octopamine-like receptors. These were used as queries to search via BLASTn in the recently released *V. destructor* genome (Techer et al. 2019). Thus, contigs c139848_g8_i1 (1227 bp), and c143491_g6_i5 (2547 bp), mapped to the locus LOC111253729, annotated as a G-protein couple receptor (XP_022669321.1), and to the locus LOC111251882, annotated as an octopamine receptor beta-2R-like (XP_022664702.1), respectively. Since in *Rhipicephalus microplus*, a tyramine receptor (GenBank accession number CAA09335) was previously associated with resistance to amitraz (Kumar 2019), the homologous gene was searched in the *V. destructor’s* genome. The locus LOC111254088, annotated as octopamine-like receptor in the *V. destructor* database, showed the highest identity with the gene encoding the CAA09335 protein from *R. microplus*. Phylogenetic analysis was then conducted with these proteins and with others, annotated as octopamine receptors, from several arthropod species (Table S2). The phylogenetic tree obtained from the alignment of 47 proteins clustered into three main groups, consisting of α_2_-adrenergic-like octopamine receptors (Octα_2_Rs), β-adrenergic-like octopamine receptors (OctβRs), and type 1 Tyramine receptors (TAR1). The branch corresponding to OctβRs included three classes of receptors: Octβ_1_R, Octβ_2_R, and Octβ_3_R. Regarding the proteins from *V. destructor* in the alignment, XP_022669321 grouped with Octα_2_Rs; XP_022664702 is included in the branch corresponding the Octβ_2_Rs, and XP_022670329 is related with TAR1s (Fig. 1). From this analysis, we called XP_022669321, XP_022664702, and XP_022670329 proteins, as Vd_Octα_2_R, Vd_Octβ_2_R, and Vd_TAR1, respectively.

**Figure 1.**
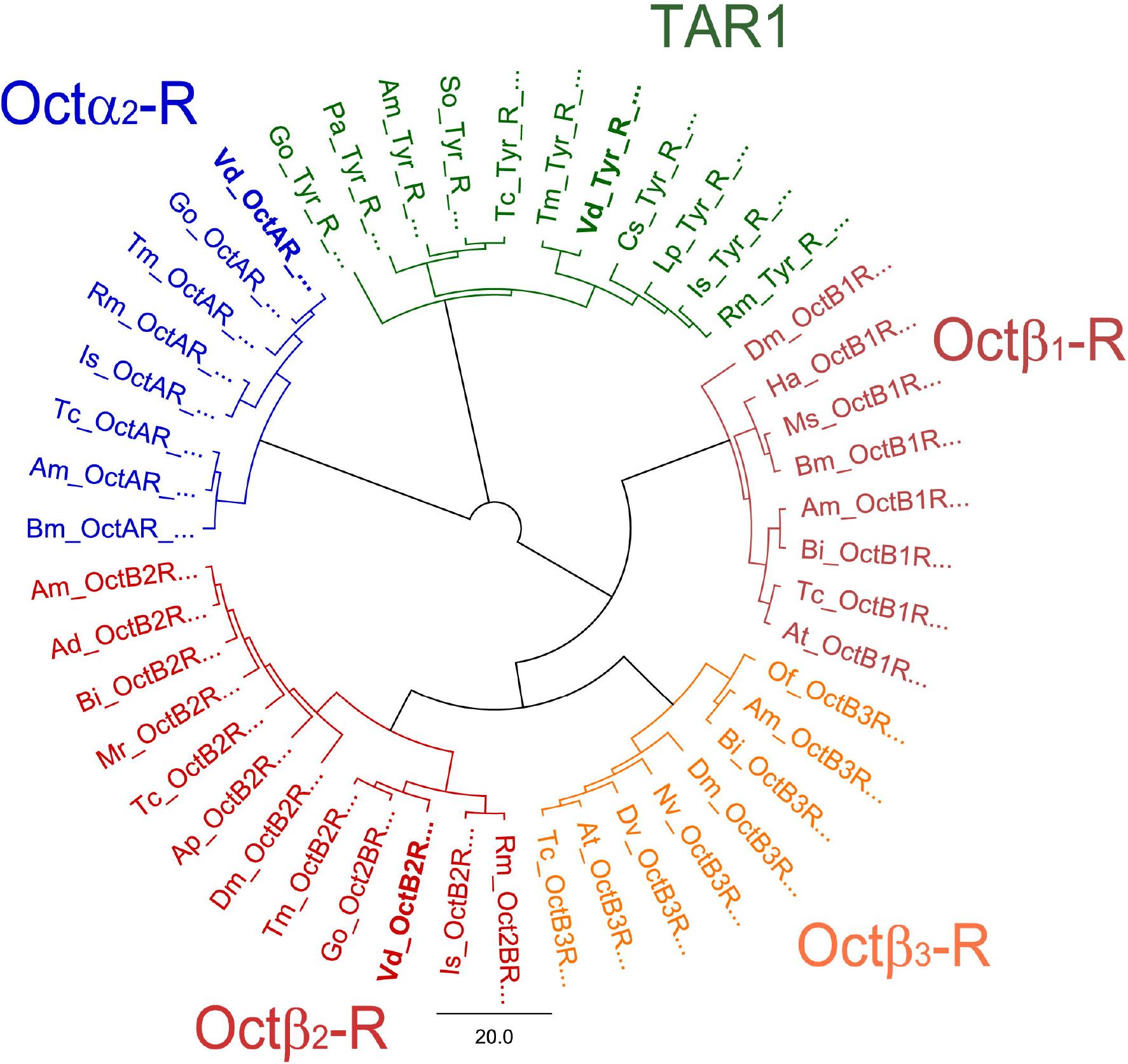
Phylogenetic tree of octopamine receptors across several species of arthropods. Neighbor-Joining tree was constructed in Geneious 9.1.8. Octα_2_R: α_2_-adrenergic-like octopamine receptor; Octβ_1_R: octopamine β_1_ receptor; Octβ_2_R: octopamine β_2_ receptor; Octβ_3_R: octopamine β_3_ receptor; TAR1: type 1 tyramine receptor. Ac: *Acyrthosiphon pisum*; Ae: *Aethina tumida*; Am: *Apis dorsata*; Am; *Apis mellifera*; Bi: *Bombus impatiens*; Bm: *Bombyx mori*; Cs: *Centruroides sculpturatus*; Dv: *Diabrotica virgifera virgifera*; Dm: *Drosophila melanogaster*; Go: *Galendromus occidentalis*; Ha: *Helicoverpa armigera*; Is: *Ixodes scapularis*; Lp: *Limulus polyphemus*; Ms: *Manduca sexta*; Mr: *Megachile rotundata*; Nv: *Nicrophorus vespilloides*; Of: *Ostrinia furnacalis*; Pa: *Periplaneta americana*; Rm: *Rhipicephalus microplus*; Tc: *Tribollium castaneum*; Tm: *Tropilaelaps mercedesae*; Vd: *Varroa destructor*. The GenBank accession numbers of the receptor sequences in the tree are listed in Table S2.

### Vd_Octα_2_R protein

Vd_Octα_2_R encoded for a 532 amino acids protein. When Vd_Octα_2_R was aligned to other α-adrenergic-like octopamine receptors, a high degree of conservation was observed among species, mainly in the regions corresponding to the predicted seven α-helices of the proteins’ tertiary structure (Fig. 2A). The percentage of identity between Vd_Octα_2_R and the α-receptors from other acari species was 82 % for *Galendromus occidentalis*, 68 % for *Tropilaelaps mercedesae*, 52 % for *R. microplus* and 50 % for *Ixodes scapularis*. Conserved motifs common to GPCR were found in α-helices III, VI, VII, and the C-terminus (Fig. 2A).

**Figure 2.**
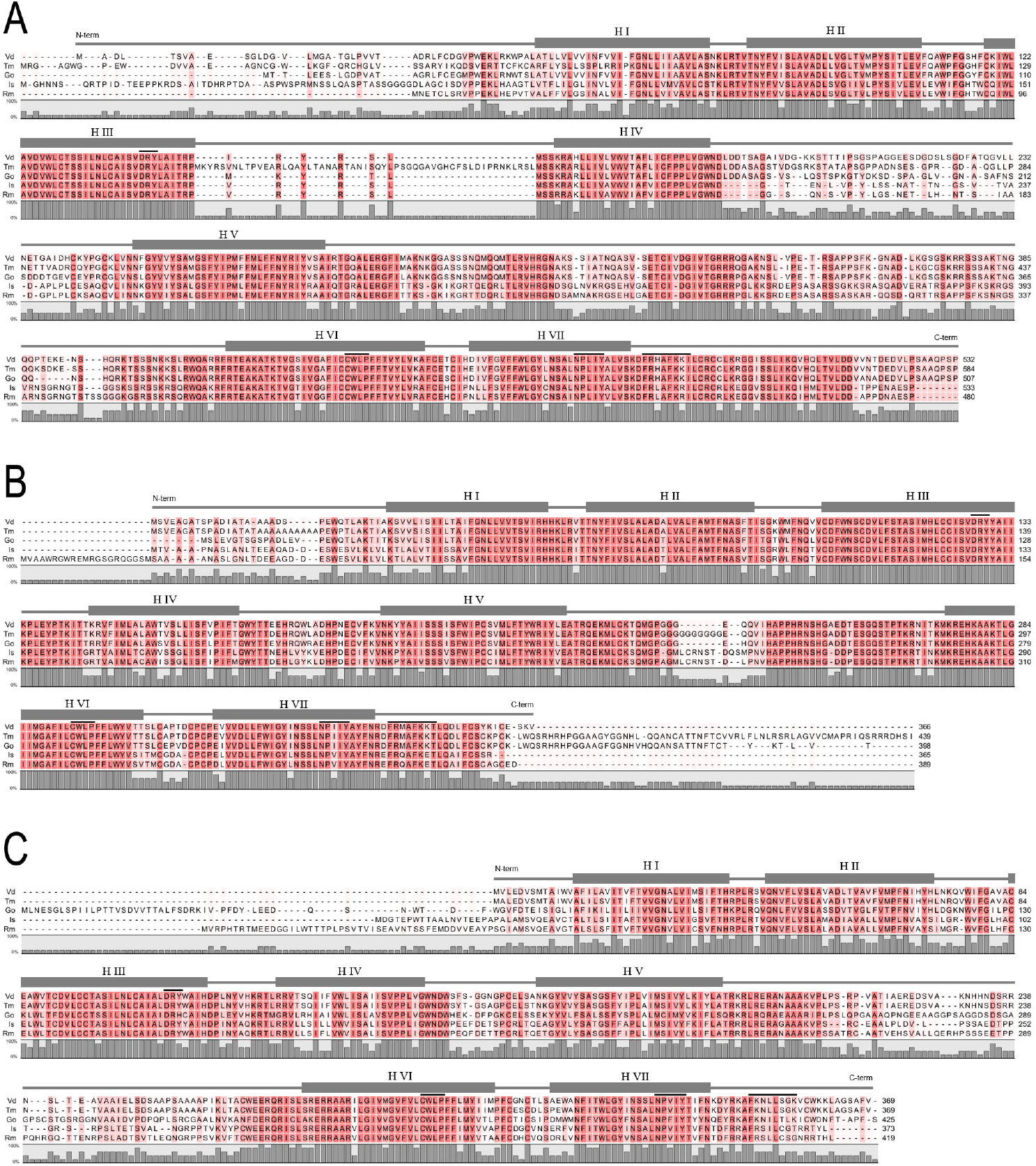
Multiple sequence alignment of OctαRs (A), OctβRs(B), and TAR1 (C) from diverse acari species. The shaded sequences highlight the amino acid identity level. The seven α-helices are represented as grey rectangles and numbered as H I-VII. GPCR conserved motifs in helix III (D[E]RY), helix VI (CWxP), helix VII (NP[L/I]IY), and C-terminus (F[R/K]xx[F/L]xxx) are indicated by bars. Vd: *Varroa destructor*; Tm: *Tropilaelaps mercedesae*; Go: *Galendromus occidentalis*; Is: *Ixodes scapularis*; Rm: *Rhipicephalus microplus*.

### Vd_Octβ_2_R protein

Vd_Octβ_2_R encoded for a 439 amino acids protein. The percentage of identity between Vd_Octβ_2_R and the β-adrenergic-like octopamine receptor from other closely related acari species in the cladogram were 83 % for *T. mercedesae*, 79 % for *G. occidentalis*, 68 % for *R. microplus* and 65 % for *I. scapularis*. Multiple sequence alignment of OctβRs from these species showed that, as in Vd_Octα_2_R, Vd_Octβ_2_R contained highly conserved regions corresponding to the seven α-helices typical of GPCR (Fig. 2B). The modelling of the Vd_Octβ_2_R three-dimensional structure, obtained with I-TASSER online server, showed the common structure described in GPCRs: seven transmembrane (TM) helical bundle connected by three extracellular loops (EL) and three intracellular loops (IL) (Fig. 3A). The N-terminus of the protein was at the extracellular side and the C-terminus was located intracellularly. In this structure, the ligand-pocket would be close to the extracellular region and surrounded by the transmembrane helical domain (Marsh 2015). The molecular simulation of transmembrane regions using Phobius software predicted which residues were “buried” into the membrane or exposed to intracellular or extracellular regions (Fig. 4). Other features characterizing OctβR were also found in Vd_Octβ_2_R (Fig. 4). The receptor had two highly conserved cysteine residues in TM3 and EL2 which form a disulphide bond, which is important for stabilizing the conformation of the extracellular region and shaping the entrance to the ligand-binding pocket (Rader et al. 2004). Three motifs of amino acids involved in molecular switches in GPCRs during activation were also found in Vd_Octβ_2_R: i) the D[E]RY motif in helix III, which often forms a so-called “ionic lock”. The ionic lock was suggested as a characteristic of the inactive conformation of GPCRs, blocking the G-protein binding at the cytoplasmic region; ii) the CWxP motif observed in α-helix VI, considered as one of the micro-switches that have substantially different conformations in the active state versus the inactive state of the receptor; iii) the NP(L/I)IY motif in helix VII, involved in a permanent rotameric change (Filipek 2019) (Fig. 2B and Fig. 4). As in most of the GPCR structures, the C-terminus contains a 3-4 turn α-helix, α-helix VIII, that runs parallel to the membrane and is characterized by a common (F[R/K]xx[F/L]xxx) amphiphilic motif (Zhang et al. 2015). Putative amino acids involved in octopamine binding are extended through a 222 amino acids region between W106 and Y327.

**Figure 3.**
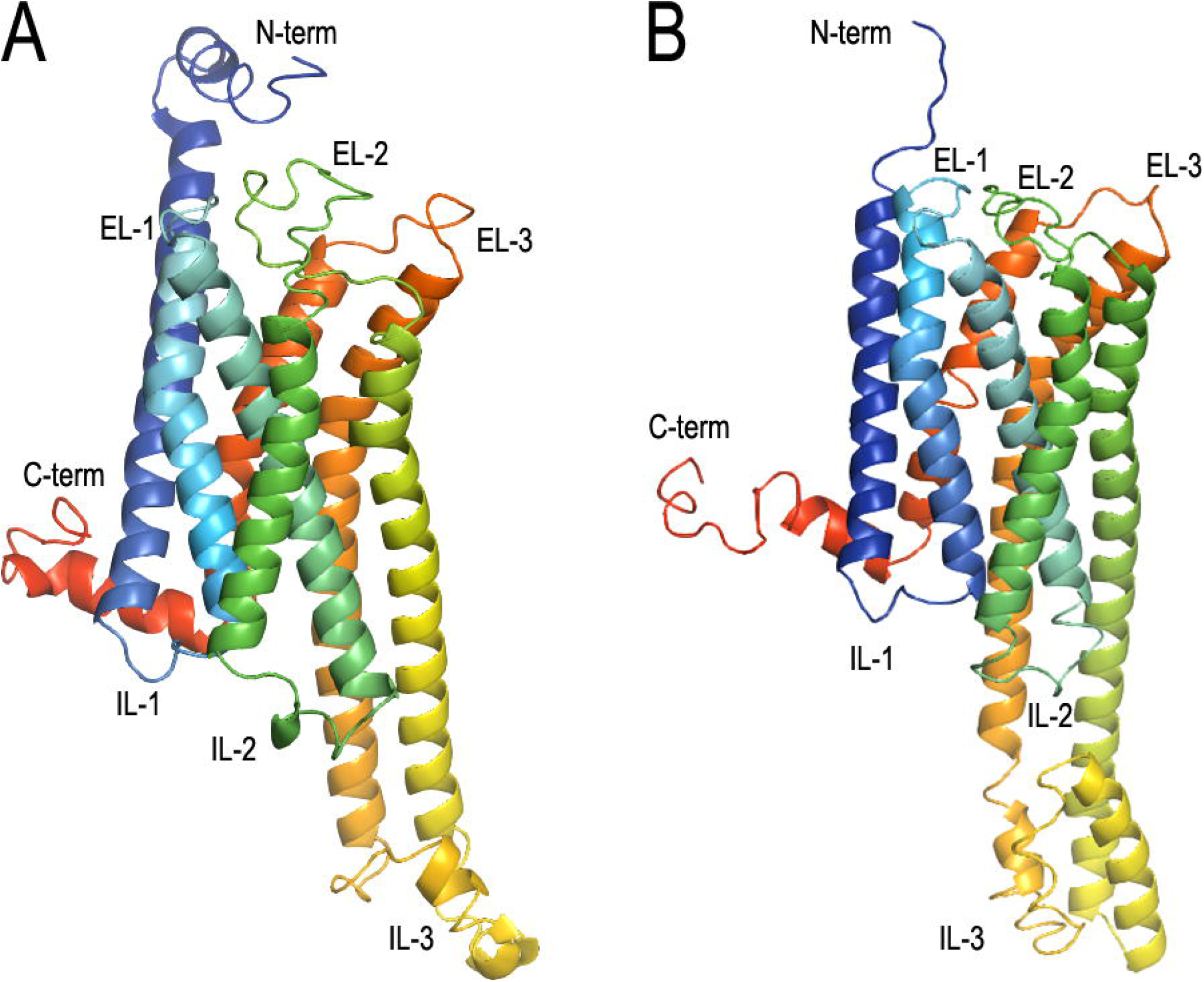
Three-dimensional structure of Vd_Octβ_2_R (A) and Vd_TAR1 (B), obtained by modelling with I-TASSER (Yang and Zhang 2015). The receptors are showed as ribbon representation in rainbow colouring (N-terminus, blue; C-terminus, red). The seven α-helices are connected by three extracellular loops (EL1-3) and three intracellular loops (IL1-3).

**Figure 4.**
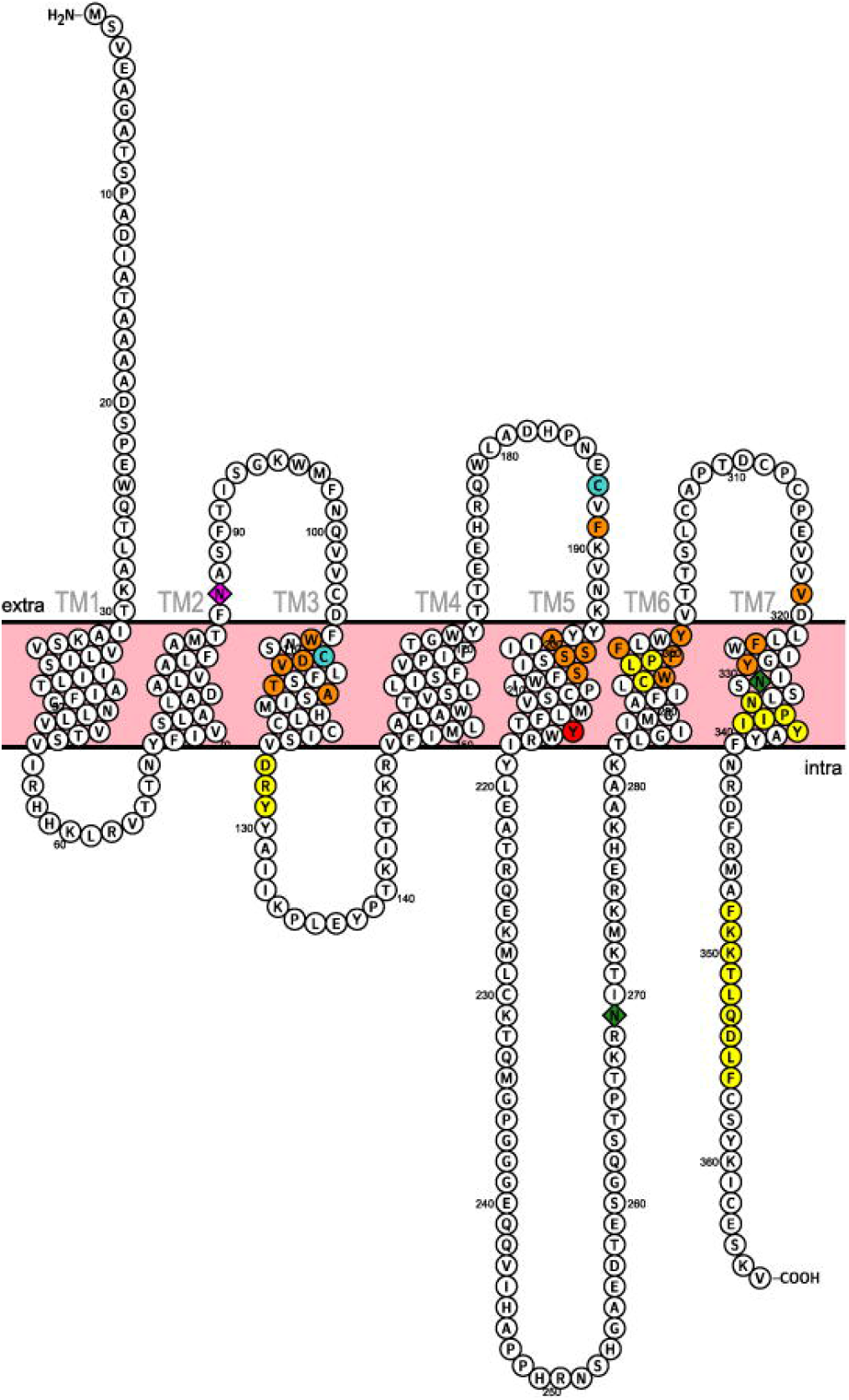
Snake plot of Vd_Octβ_2_R with transmembrane domains predicted with Phobius (Kall et al. 2004). N87S mutation (magenta); Y215 mutation (red); putative N-glycosylation residues (diamond); GPCR conserved motifs (yellow); putative disulphide bond residues (blue); predicted ligand binding residues (orange).

### Vd_TAR1 protein

Vd_TAR1 encoded for a 369 amino acids protein. Vd_TAR1 was aligned to the acari tyramine receptors more similar to the tyramine receptor of *R. microplus* (CAA09335), in which mutations associated with resistance to amitraz have been described (Kumar 2019). As with Vd_Octα_2_R and Vd_Octβ_2_R, the regions corresponding to the predicted seven helices in the tertiary structure of the proteins are conserved among species (Fig. 2C). The modelling of the three-dimensional structure Vd_TAR1 also showed the described structure for GPCR: seven hydrophobic transmembrane domains and six hydrophilic loops (Fig. 3B). Like in other TAR1 receptors, the third intracellular loop of Vd_TAR1 is longer than that in Octβ_2_Rs. The percentage of identity between Vd_TAR1 and the tyramine receptors from other acari species is 94 % for *T. mercedesae*, 61 % for *R. microplus* and for *I. scapularis*, and 56 % for *G. occidentalis*.

### *Vd_octα*_*2*_*r, Vd_octβ*_2_*r* and *Vd_tar1* genes

The cDNA of *Vd_octα*_*2*_*r, Vd_octβ*_*2*_*r* and *Vd_tar1* were obtained by RT-PCR, using as template the same RNA samples used for transcriptomics. Sequencing of the ORFs showed a full identity of these cDNAs with XM_022813586, XM_022808967 and XM_022814594, corresponding to the mRNA of Vd_Octα_2_R, Vd_Octβ_2_R, and Vd_TAR1, respectively.

The ORF of *Vd_octα*_2_*r* has a length of 1,599 bp, and the full gene is 155,562 bp long. The *Vd_octα*_2_*r* gene comprises nine exons and eight introns (Fig. 5A). The 5’UTR is extended along Exon 1, Exon 2 and Exon 3. The start codon (position 3,746 at the mRNA) is sited in Exon 4. The stop codon (position 5,344 at the mRNA) and the 3’UTR are in the Exon 9, the largest exon. The length of all the exons and introns of *Vd_octα*_2_*r* is shown in Fig. 5A.

**Figure 5.**
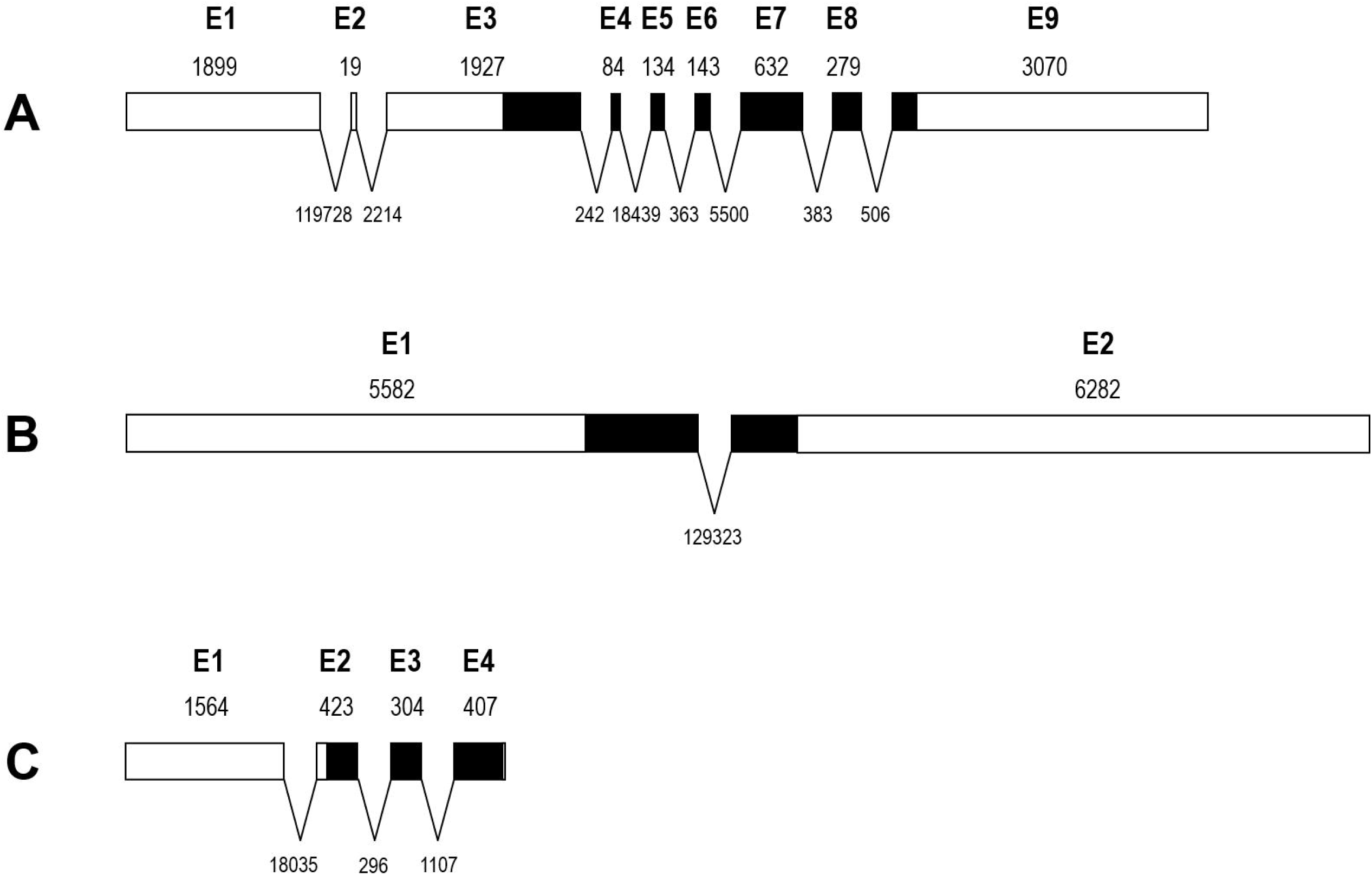
Schematic diagram of *octα*_*2*_*r* (A), *octβ*_*2*_*r* (B), and *tar1* (C) exon-intron gene structure. Coding sequence (CDS) are shown in black. Lengths are represented in bp.

The lengths of *Vd_octβ*_2_*r* ORF, mRNA, and full gene sequences are 1,101 bp, 11,863 bp, and 141,186 bp, respectively. The *Vd_octβ*_2_*r* gene comprises two exons and one intron (Fig. 5B). Exon 1 contains the 5’UTR and the start codon (position 4,507 at the mRNA), and Exon 2 contains the stop codon (position 5,607 at the mRNA) and the 3’UTR. Between Exon 1 and Exon 2 there is a long intron of 129,323 bp (Fig. 5B).

The *Vd_tar1* gene has a length of 22,226 bp, transcribed into an mRNA of 2,788 bp in which an ORF of 1,110 bp is found. The *Vd_tar1* gene comprises 4 exons and 3 introns (Fig. 5C). The 5’UTR is extended along Exon 1 and Exon 2. The start codon (position 1,162 at the mRNA) is sited in Exon 2. The stop codon (position 2,769 at the mRNA) and the 3’UTR are in the Exon 4. The length of all the exons and introns of *Vd_tar1* is shown in Fig. 5C.

### *Vd_octβ*_*2*_*r* and *Vd_tar1* sequences in *V. destructor* mites susceptible to amitraz

Total RNA was isolated from pools of five to ten *V. destructor* adult females collected in Iran, New Zealand, Spain and the UK between 2016 and 2019 from colonies without amitraz treatment. As mutations associated with the resistance to amitraz has been described in OctβR and TAR1 receptors, RNA from these susceptible mites was reverse transcribed into cDNA to amplify the full length of *Vd_octβ*_*2*_*r* and *Vd_tar1* ORFs. The sequencing of *Vd_octβ*_*2*_*r* and *Vd_tar1* ORFs of mites from these countries showed identical sequences to those previously identified as wild-type in this paper (XM_022813586 and XM_022814594, respectively).

### *Vd_octβ*_*2*_*r* N87S mutation

We identified a single point mutation in *Vd_octβ*_*2*_*r* gene (substitution of A to G at nucleotide 260 of the ORF) in mites extracted alive from the brood, right after finishing the treatment with amitraz, in colonies of apiary DTRA (Isère department, France), that reported failure of this treatment. This mutation results in an asparagine (AAT) to serine (AGT) substitution at position 87 of the Vd_Octβ_2_R protein (N87S) (Fig. 6A). To validate this result, total DNA was isolated from 24 individual mites collected in 3 colonies from the same apiary. The genomic region comprising the mutation was amplified and sequenced. All sequenced mites showed the N87S mutation (Table 1). The same analysis was carried out with mites from the apiaries MAP (Loire department) and MHRA (Isère department), where the treatment with amitraz also failed. The mutation was present in 75 % of the mites from MAP apiary, and in 71 % of the mites from the MHRA apiary (Table 1). In colonies MM16 and J11 (both located in apiaries at Puy-de-Dôme department) the mutation N87S was detected in 77 and 57 % of the mites, respectively (Table 1). Further analysis showed that the efficacy of amitraz treatment was 92 % in colony MM16 and 77 % in J11. The occurrence of this mutation was also studied in three apiaries from nearby departments in which amitraz was not used the year before sampling. Apiaries VB (Alpes-de-Haute-Provence department), AmA (Maine-et-Loire department) and DE (Drôme) were all treated with oxalic acid. None of the mites from VB and AmA carried the mutation N87S while 26 % of the mites from DE were mutants (Table 1). Altogether, these data show circumstantial evidence that there is an association between the mutation N87S and amitraz treatment failure.

**Table 1.** Frequency of the N87S mutation in the samples collected from several French departments.

| Colony | Department | n | Last treatment | Collection | % N87S mutation |
| --- | --- | --- | --- | --- | --- |
| DTRA | Isère (38) | 24 | Amitraz | POST-Treatment | 100 % |
| MAP | Loire (42) | 24 | Amitraz | POST-Treatment | 75 % |
| MHRA | Isère (38) | 24 | Amitraz | POST-Treatment | 71 % |
| J11 | Puy-de-Dôme (63) | 23 | Amitraz | POST-Treatment | 74 % |
| MM16 | Puy-de-Dôme (63) | 24 | Amitraz | POST-Treatment | 58 % |
| VBA | Alpes-de-Haute-Provence (04) | 20 | Oxalic acid | PRE-Treatment | 0 % |
| AmA | Maine-et-Loire (49) | 24 | Oxalic acid | PRE-Treatment | 0 % |
| DE | Drôme (26) | 19 | Oxalic acid | PRE-Treatment | 26 % |

**Figure 6.**
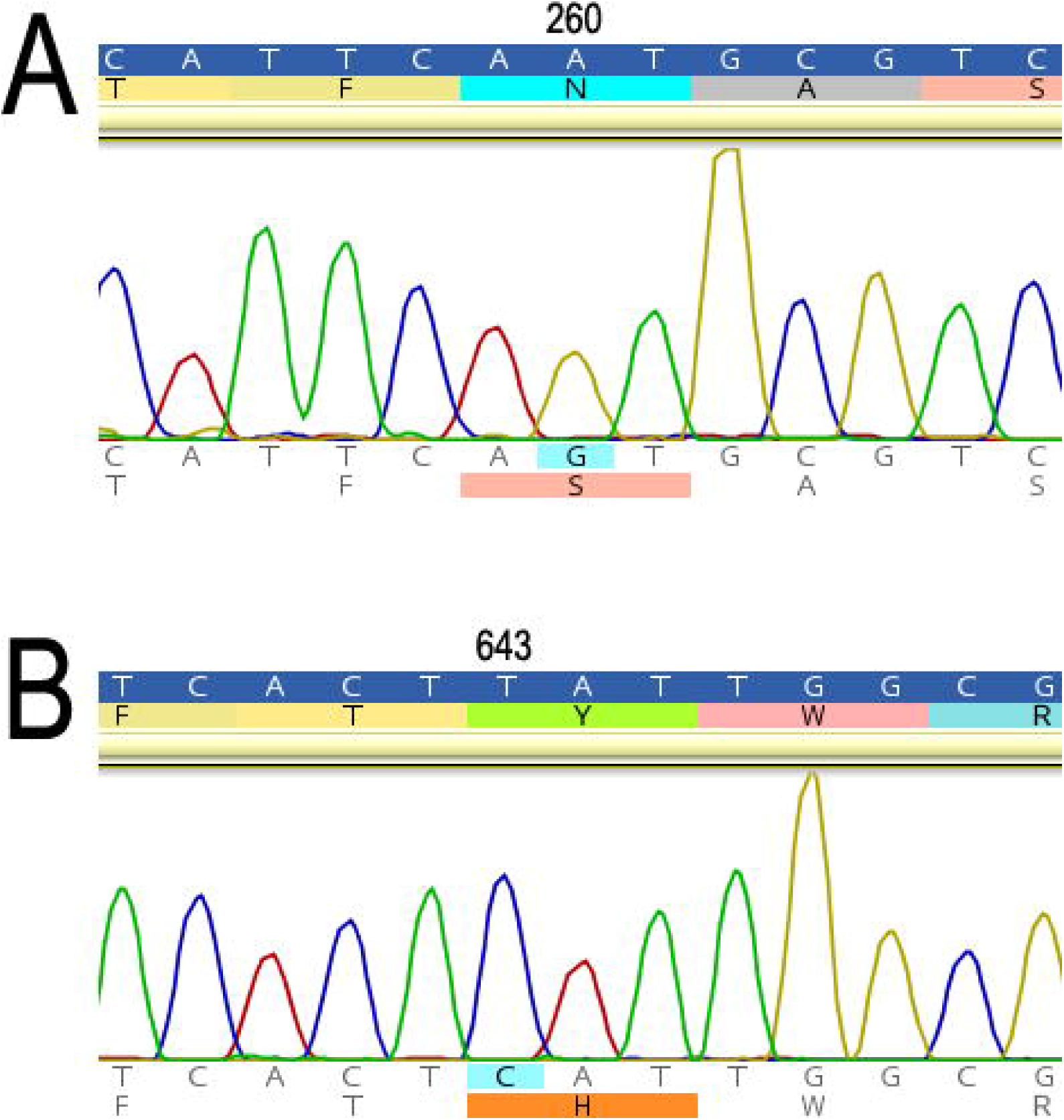
Electropherograms showing the mutations in the sequence of Vd_Octβ_2_R. The substitution of A by G at position 260 of the ORF results in the N87S mutation (A), whereas the substitution of T by C at position 643 results in the Y215H mutation (B).

The ORF of *Vd_tar1* was also sequenced in pools of mites collected from all French apiaries analysed in this study. None of the analysed mites showed any change in the sequence when compared with the wild-type *Vd_tar1*.

### Y215H mutation

In the U.S., a state apiary inspector reported the failure of the amitraz treatment in some colonies from New Jersey in 2018 (Styles, Personal communication). The *Vd_octβ*_*2*_*r* and *Vd_tar1* gene sequences were examined in mites collected from four of these colonies. No mutations were detected in *Vd_tar1* gene and the mutation N87S, identified in French samples, was also not detected. However, a new single point mutation was identified in the *Vd_octβ*_*2*_*r* gene from mites collected from the four colonies. The substitution of T to C at position 643 of the ORF results in a tyrosine (TAT) to histidine (CAT) substitution at position 215 of the Vd_Octβ_2_R protein (Y215H) (Fig. 6B). This mutation was detected in 50 to 96 % of the mites sequenced from these colonies (Fig. 7, Table S1).

**Figure 7.**
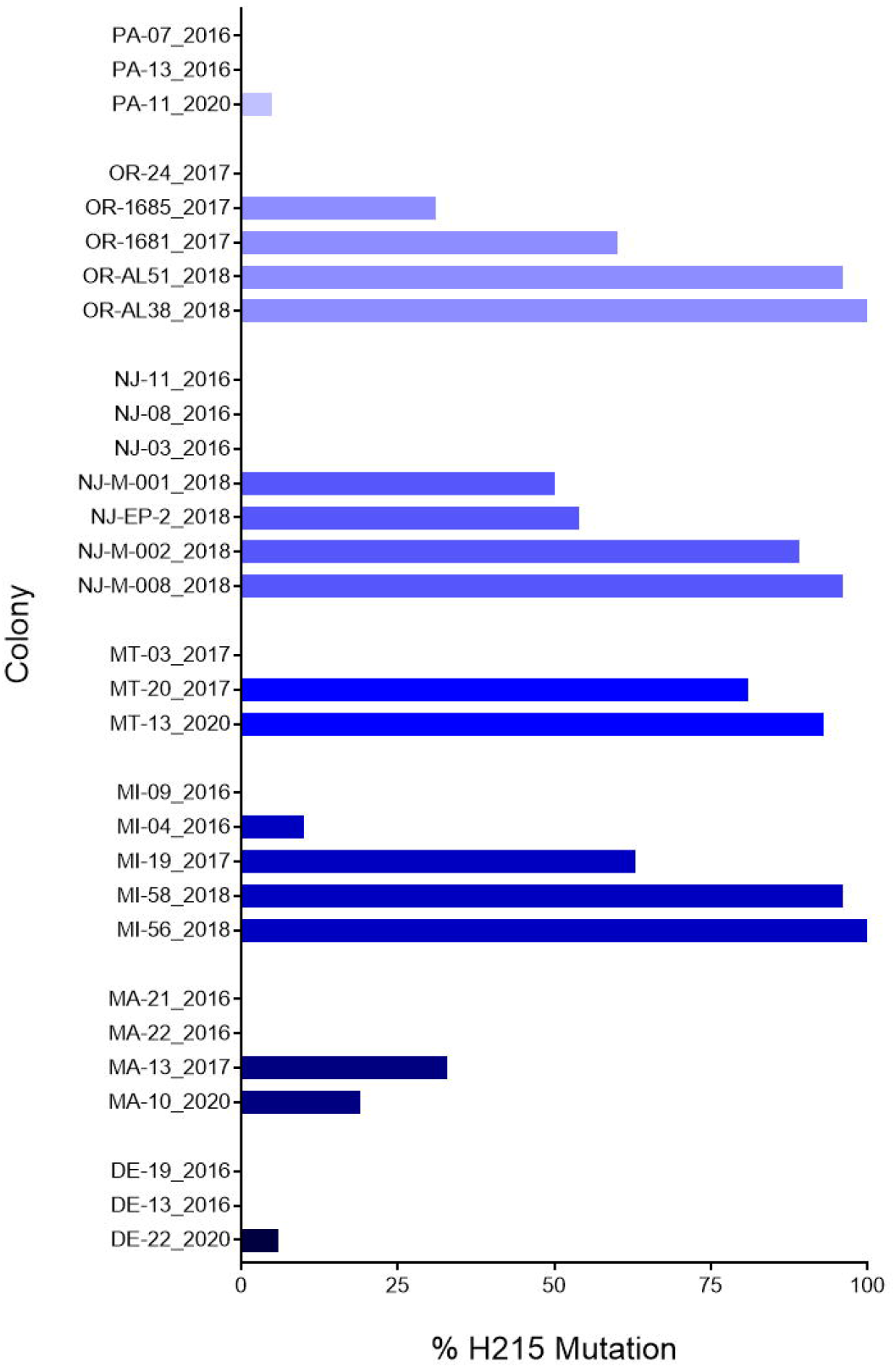
Timeline of the Y215H mutation incidence in colonies from U.S. The name of the colonies in the Y axe shows the state and the year of sample collection. DE: Delaware; MA: Massachusetts; MI: Michigan; MT: Montana; NJ: New Jersey; OR: Oregon; PA: Pennsylvania. More detailed information can be found in Table S1.

In order to gather data regarding the presence of the mutation Y215H in New Jersey from previous years, mites collected in 2016 from different colonies in this state were also sequenced. We did not detect this mutation in any of the colonies analysed (Fig. 7, Table S1).

Since the presence of the Y215H mutation seemed related with the reduced susceptibility to amitraz, we analyzed mite samples obtained from a BIP project evaluating the efficacy to Apivar^®^ in Oregon and Michigan in 2018. These trials were suggestive of amitraz treatment failure (Nathalie Steinhauer, personal communication). Samples of phoretic *V. destructor* mites were collected from bees sampled from colonies while being treated with Apivar^®^. The Y215H mutation was detected in 88 and 96 % of the mites from the two colonies we examined that were treated with Apivar^®^ in Oregon, and in the 94 and 90 % of the mites from the two colonies treated with Apivar^®^ in Michigan (Fig. 7, Table S1). Colonies from the same apiaries but treated with thymol instead of Apivar^®^ were also analyzed. The Y215H mutation was present in 96 and 100 % of the mites collected in the two colonies from Oregon and the same frequencies were also recorded in the two colonies from Michigan (Table S1). On the other hand, mites collected from these two states before 2018 (Millán-Leiva et al. 2021a) were also sequenced. The mutation was found but at much lower frequency, suggesting that the mutation is a relatively recent event (Fig. 7, Table S1).

To estimate when the mutation first evolved in the U.S. population, we compared the presence of Y215H in samples collected in 2020 with samples collected in previous years in several U.S. states (Millán-Leiva et al. 2021a). Results from Delaware, Massachusetts, Montana and Pennsylvania showed that the mutation was practically non-existent in 2016 but its incidence has increased since (Fig. 7, Table S1).

### Diagnostic assay

Two high throughput allelic discrimination assays based on TaqMan^®^ were developed to enable rapid and accurate genotyping of N87S and Y215H mutations in individual mites. For each real-time PCR assay, we designed two fluorescent labelled probes to discriminate between wild-type and mutant alleles. The probes selective for N87 or Y215 wild-type alleles were labelled with VIC^®^ while the others, selective for S87 or H215 alleles, were labelled with 6FAM™. Therefore, an increase in VIC^®^ fluorescence indicates the presence of the wild-type allele, while an increase in 6FAM™ fluorescence indicates the presence of the mutant allele. An intermediate increase in the fluorescence of both dyes indicates that the mite is heterozygous for the mutation. Twenty-four mites in which the nucleotide at each of the mutation sites of *Vd_octβ*_*2*_*r* was known by previous sequencing were genotyped by TaqMan^®^. The results showed a perfect correlation between data from sequencing and genotyping. Genotyped mites were either homozygous for the wild-type allele (N87 or Y215), the mutant allele (S87 or H215), or heterozygous for each mutation (Fig. 8).

**Figure 8.**
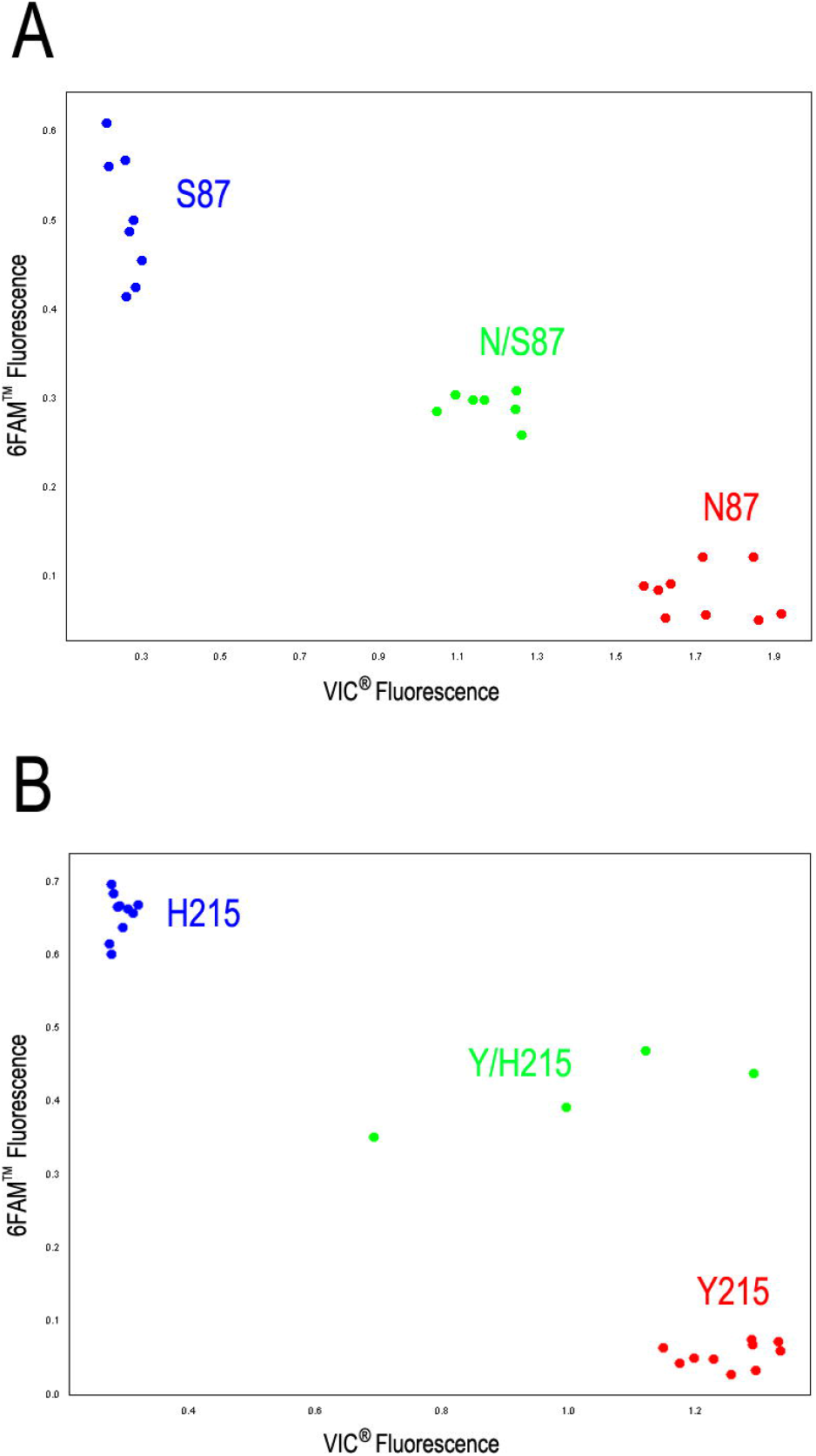
Real-time TaqMan^®^ detection of the N87S (A) and Y215H (B) mutations in Vd_Octβ_2_R. In the scatter plots of VIC^®^ and 6FAM™ fluorescence, each dot represents an individual mite. SS homozygotes (N87 or Y215 allele) in red; RS heterozygotes in green; RR homozygotes (S87 or H215 allele) in blue.

## DISCUSSION AND CONCLUSION

Here we identified two amino acid substitutions, located in the β-adrenergic octopamine receptor of *V. destructor*, that seem to be associated with field treatment failures using amitraz in samples collected in France and the U.S. Our data also show circumstantial evidence of an independent evolution of resistance in both locations.

Amitraz is a formamidine that has been widely used as an acaricide since its discovery back in 1972 (Harrison et al. 1972). Nowadays, it is one of the main alternatives for controlling varroosis worldwide. This compound mimics the action of the neurotransmitters octopamine and tyramine and blocks their receptors (Hollingworth and Lund 1982). Therefore, it is likely that modifications in key sites of the octopamine or tyramine receptors would be associated with the treatment failures reported by beekeepers after treatments with amitraz-based acaricides.

A joint analysis of transcriptomic (BioProject ID PRJNA531374) and genomic data (Techer et al. 2019), alongside with data available in public databases, allowed the characterization of proteins from three different classes of receptors in this mite: an α-adrenergic-like octopamine receptor (Vd_Octα_2_R), a β-adrenergic-like octopamine receptor (Vd_Octβ_2_R) and a tyramine type 1 receptor (Vd_TAR1). A more in-depth *in silico* study of the secondary and tertiary structures of these proteins showed that they have structural features typical of the superfamily of G-protein coupled receptors, such as the seven transmembrane domains and the classic distribution of extracellular and intracellular loops (Finetti et al. 2021). Moreover, the occurrence of highly conserved residues and several sequence motifs common to α- and β-adrenergic octopamine receptors in Vd_OctαR and Vd_OctβR, confirmed the correct identification and classification of these proteins as octopamine receptors in *V. destructor*. It was once thought that amitraz only interacts with octopamine receptors (OAR). However, during that time, tyramine type 1 receptors have been wrongly classified as OAR (Chen et al. 2007). Later, this receptor was classified as Oct/TyrR (Baron et al. 2015) and recently, tyramine type 1 receptor was finally classified as TAR (Farooqui 2012; Finetti et al. 2021). However, as this is a recent change in the classification, it is still not updated in public databases, that maintain erroneous annotations, leading to confusion when trying to identify and classify this family of receptors. This is the case of *V. destructor*, in which Vd_TAR1 (XP_02270329) is described as octopamine receptor-like, actually being a tyramine receptor, as we have thoroughly described in this study.

Resistance to amitraz in Varroa have been reported in populations from different locations around the world, such as the U.S. (Elzen et al. 1999; Elzen et al. 2000; Rinkevich 2020), Mexico (Rodríguez-Dehaibes et al. 2005), Argentina (Maggi et al. 2010), the Czech Republic (Kamler et al. 2016) and France (Almecija et al. 2020). In addition to these publications, anecdotal reports of reduced amitraz efficacy are widely discussed among beekeepers (Rinkevich 2020). However, until now, the mechanism causing this lack of efficacy was unknown.

The mechanism of resistance to amitraz has been thoroughly studied in the cattle tick *R. microplus* (Baxter and Barker 1999; Chen et al. 2007; Corley et al. 2013; Baron et al. 2015; Koh-Tan et al. 2016; Jonsson et al. 2018). In this species, the resistance detected in the field has been associated with polymorphisms in the octopamine and tyramine receptors, suggesting that target site insensitivity is the most common mechanism of resistance to amitraz. Chen et al. (2007) found two amino acid substitutions (T8P and L22S) in the tyramine receptor gene that were only present in American strains highly resistant to amitraz. Further analysis by Baron et al. (2015) supported the association of these two SNPs with the resistance in field samples collected in South Africa. However, previous analysis of the same gene with samples collected in Australia did not find any SNPs differentiating susceptible from resistant strains (Baxter and Barker 1999). In an attempt to address this issue, Corley et al. (2013) widen the scope of the analysis to other octopamine receptors using the same amitraz-resistant Ultimo strain analysed by Baxter and Barker. They found an increased frequency of the mutation I61F in the β-adrenergic octopamine receptor (RmBAOR) providing circumstantial support for associating this mutation with the resistance to amitraz in the Ultimo strain. Supporting this association, an I45F mutant of *Bombyx mori* OAR2 (equivalent to I61F in RmBAOR) showed reduced sensitivity to the amitraz metabolite DPMF (N^2^-(2,4-Dimethylphenyl)-N^1^ methyformamidine) in HEK-293 cells (Takata et al. 2020). In a different study, cell lines derived from acaricide-resistant *R. microplus* colonies from Colombia contained a 36 bp duplication in the RmBAOR gene leading to a 12 amino acid insertion in the first transmembrane domain of the protein (Koh-Tan et al. 2016). Further analyses of resistant *R. microplus* from Brazil, Mexico, Australia, Thailand and South Africa supported the association of I61F with the resistance, but also described novel SPNs in the RmBAOR associated with amitraz resistance in specific populations (Jonsson et al. 2018).

To date, there is no reported association between mutations in α-adrenergic octopamine receptors and resistance to amitraz. Therefore, we analysed Vd_TAR1 and Vd_Octβ_2_R, the receptors of *V. destructor* phylogenetically closer to those of *R. microplus* reporting polymorphisms associated with amitraz resistance. None of the mutations described in *R. microplus* were found in the *V. destructor* samples analysed in this study. However, we did identify two novel non-synonymous substitutions in the *Vd_octβ*_2_*r* gene with a differential geographical distribution. A substitution of asparagine 87 to serine (N87S) associated with treatment failures in France, and a substitution of tyrosine 215 to histidine (Y215H) in samples collected across the U.S. from colonies reporting low amitraz efficacy. None of the samples analysed in this study were collected as part of a structured sampling strategy designed to elucidate the mechanism of resistance to amitraz. Rather, most of them were part of projects, experiments or surveys conducted to validate previous reports of treatment failures. After a careful case-by-case analysis of the sampling and treatment history, it is possible to draw conclusions on whether these mutations are associated with the resistance to amitraz. In the case of samples collected in France, when the sampling was conducted after finishing the treatment with amitraz (Table 1), a significant number of mites were mutants for N87S (always above 50 %), showing an association with the efficacy observed in the field. On the other hand, the samples collected from colonies not exposed to amitraz at least the year before the sample collection were mostly wild-type. This suggests that amitraz is exerting a significant selection pressure, favouring the prevalence of N87S mutants in the populations after an intensive treatment regime for many years. In the U.S., the samples were collected as part of different projects and screening efforts using different sampling approaches. In these cases, whenever the mites (phoretic) were collected after finishing the treatment with amitraz (NJ-M-NJ-M-001, NJ-M-002, NJ-M-008, NJ-EP-2) or when the treatment was still ongoing (OR-AV01, OR-AV02, MI-22, MI-33), the frequency of mutants was very high (Table S1), indicating an association between the presence of the mutation Y215H and the survival after exposure. However, the samples collected from other colonies (OR-AL38, OR-AL51, MI-56, MI-58), taking part in the same field assay in Oregon and Michigan but treated with thymol, also showed a high frequency of mutant mites. This may be explained considering that amitraz has been used intensively for long time in these locations. Thus, given the high movement of mites within apiaries (Kulhanek et al. 2021), it is possible that a significant part of the population was already mutant before starting the field trials in 2018. The historical data gathered after the analysis of samples collected in 2016 and 2017 also supports this idea. Our data show that the mutation was nearly absent in the samples collected in several states in 2016, with only one sample with mutants in Michigan (MI-09). Yet, in 2017, although some samples were still completely wild-type, many of them show that the mutation was present in a significant number of mites. Hence, it is reasonable to think that in 2018, following the same treatment regime with amitraz, the frequency of mutants -e.g. resistant mites-would predominate (Table S1).

The joint analysis of the data also suggests that the resistance have evolved independently at both locations. The mutation N87S was detected only in mites collected in France while Y215H was detected only in the mites collected in the U.S. This result is yet another example of the capacity of this species to evolve resistance to the same acaricide via multiple independent pathways. This was already described for the resistance to pyrethroids based-acaricides. In Europe mites carry mostly the mutation L925V in the VGSC, while those from the U.S. carry the mutations L925M and L925I (González-Cabrera et al. 2013; González-Cabrera et al. 2016; González-Cabrera et al. 2018; Millán-Leiva et al. 2021a). A more recent study also evidenced that this was the result of a parallel and independent evolution process (Millán-Leiva et al. 2021b). Following the same rationale, the different mutations associated with the resistance of *R. microplus* to amitraz that evolved in different locations, in different receptor proteins and also in different residues of the same protein (Chen et al. 2007; Corley et al. 2013; Koh-Tan et al. 2016; Jonsson et al. 2018), are a very good example of the many possibilities that can be found in *V. destructor*. As we have screened a relatively small number of samples, from few locations, a larger screening effort is called for to draw a more accurate and complete picture of the situation.

A thorough *in silico* analysis of the β-octopamine receptor of *Schistocerca gregaria* showed that the nonpolar residues of the transmembrane regions are buried in the receptor core to form a hydrophobic pocket (active pocket) that is closed to the extracellular region and surrounded by the transmembrane domain (Lu et al. 2017). According to the *in silico* model, asparagine 87 is located at the end of helix II of Vd_Octβ_2_R (Fig. 3), positioned near to the residues predicted as the putative binding site for octopamine. In the N87S mutation, the mutant residue is smaller and more hydrophobic (N -0.78; S -0.18) (Eisenberg et al. 1984) than the wild-type residue and this might lead to loss of hydrogen bonds and/or disturb the correct folding of the protein. Since this mutation is in a domain that is important for the main activity of the receptor, it might somehow disturb its function. A more targeted study found out that in *Sitophilus oryzae* amitraz and octopamine might not share the same binding site, although the two sites were close to one another (Braza et al. 2019). Docking of amitraz to *S. oryzae* tyramine receptor showed eight residues of the receptor closely interacting with this ligand. One of these amino acids was Asn91, corresponding to Asn87 in *V. destructor*. When this position was examined across species, it was found that this residue was totally conserved in both, β-adrenergic octopamine and tyramine receptors (Fig. S1A). On the other hand, in α-adrenergic-like octopamine receptors, this position shows a serine residue instead of an asparagine, indicating a possible different interaction of amitraz with OctαRs in comparison with OctβRs and TAR1s. Indeed, Kita et al. (2017) showed that the potency of amitraz and its metabolite DPMF to activate *B. mori* octopamine receptors was 347- and 2274-fold higher in β-adrenergic-like octopamine receptors than in α-adrenergic□like octopamine receptors, respectively. Additionally, based on the consensus sequence for N-linked glycosylation (NXT/S), residue N87 is predicted as a putative N-glycosylation site in Vd_Octβ_2_R. N-glycosylation has been shown to be important for many GPCRs especially in correct folding, surface expression, signalling, and dimerization (Nørskov-Lauritsen and Bräuner-Osborne 2015; Patwardhan et al. 2021). Actually, it has been reported that N-glycosylation of the α_1D_-adrenergic receptor is required for correct trafficking and complete translation of a nascent, functional receptor (Janezic et al. 2020), and that the N-glycosylation of the β_2_-adrenergic receptor regulates its function by influencing receptor dimerization (Li et al. 2017). Therefore, if the asparagine at position 87 of the Vd_Octβ_2_R is indeed a N-glycosylation site, its substitution for a serine residue may affect the integrity and functionality of this receptor.

The mutation Y215H is sited in the fifth transmembrane segment of the Vd_Octβ_2_R (Fig. 4). In this case, the wild-type residue is more hydrophobic than the mutant residue (Y 0.26; H -0.4) (Eisenberg et al. 1984). After *in silico* analysis, the prediction results based on secondary structure showed a negative effect of the substitution (score +50 with SNAP2; 100 % probability of damage with PolyPhen2). The analysis of the tertiary structure of the mutant protein indicated a decrease of the stability (Reliable index: 8 with I-MUTANT), and predicted that the hydrophobic interactions, either in the core of the protein or on the surface, would be lost (HOPE). Therefore, it seems that the change from tyrosine to histidine in this domain of the protein could seriously alter the conformation of the helix and its surroundings, which can affect the interaction of the receptor with the ligand. This hypothesis is supported by the conservation of the tyrosine residue at this position of the protein among all species analysed in this study. (Fig S1B).

Amitraz exerts its acaricidal action as an agonist of octopamine. In invertebrates, octopamine acts as neurotransmitter, neuromodulator, and neurohormone, playing a fundamental role on physiological processes (Farooqui 2007). By binding to G-coupled receptors on the surface of neurons and other cells, octopamine functions as neurotransmitter affecting diverse behaviours such as excitation, aggression and egg laying (Roeder 2005). In ticks, sublethal and behaviour effects of amitraz are considered more important than lethality in the mode of action. It has been shown that amitraz causes hyperactivity, leg waving, detaching behaviour and inhibition of the reproduction (Page 2008). Therefore, the effect of amitraz goes beyond killing like a poison; it is effective by acting as a behaviour disruptor, inhibiting the mites’ ability to remain attached to the bees before killing them. This suggests that lab bioassays that only measure LD_50_ may underestimate resistance as it would express under field conditions. Thus, looking for associations between the presence of mutations and the survival of mites in colonies treated under field conditions, is perhaps a more appropriate approach to elucidate the mechanism of resistance to products that cause behavioural changes that result in death, rather than cause death directly.

Our findings supports the association of the mutations N87S and Y215H in the β-adrenergic-like octopamine receptor of *V. destructor* with the resistance to amitraz reported in the field. Future research is needed to show a causal relationship between these mutations and the evolution of resistance to amitraz, but these tests must account for the behavioural changes induced by amitraz. Moreover, data from functional analysis via electrophysiology and other approaches will help to fully characterise the interaction of amitraz with wild-type and mutant receptors.

The current status in the management of *V. destructor* shows i) a widespread resistance to pyrethroids (Kim et al. 2009; Bak et al. 2012; González-Cabrera et al. 2016; Kamler et al. 2016; González-Cabrera et al. 2018; Millán-Leiva et al. 2021a); ii) increasing cases of failures after treatments with coumaphos (Elzen and Westervelt 2002; Maggi et al. 2009; Maggi et al. 2011); iii) and the overreliance of beekeepers on amitraz (Haber et al. 2019), which may favour the evolution of resistance to this acaricide. In this scenario, monitoring the resistance to acaricidal compounds is crucial to decide whether a given treatment is likely to be successful, as well as to avoid selection pressures with treatments that can lead to an increase of mites carrying mutations conferring resistance. To help on this endeavour, we have developed high throughput allelic discrimination assays based on TaqMan^®^ for detecting N87S and Y215H mutations in the Vd_Octβ_2_R, as was previously implemented to detect mutations in the *V. destructor* VGSC associated with resistance to pyrethroids (González-Cabrera et al. 2013; González-Cabrera et al. 2016). This assay is relatively cheap, fast, robust and capable of accurately genotype individual mites in poor quality samples. Therefore, the implementation of allelic discrimination assays like those described in this study will be especially suited towards determining the distribution and frequency of mutations associated to resistances in local Varroa populations. This information would be very valuable for designing a more rational control of Varroa, selecting each time the best acaricide for their apiaries.

## Acknowledgements

The authors thank the Bee Informed Partnership (thanks Dr. Nathalie Steinhauer, and Project Apis m), beekeepers and beekeepers’ associations from different countries for providing the mite samples used in this study. Thanks to Klemens Krieger for his support and personal implication at the beginning of this research effort. A previous version of this manuscript was revised by Frank Rinkevich (USDA, Baton Rouge, Louisiana) his comments and suggestions were used to improve the final version.

