## Supplementary figures and images for "Resistance to amitraz in the parasitic honey bee mite *Varroa destructor* is associated with mutations in the β−adrenergic-like octopamine receptor"

### Fig. S1

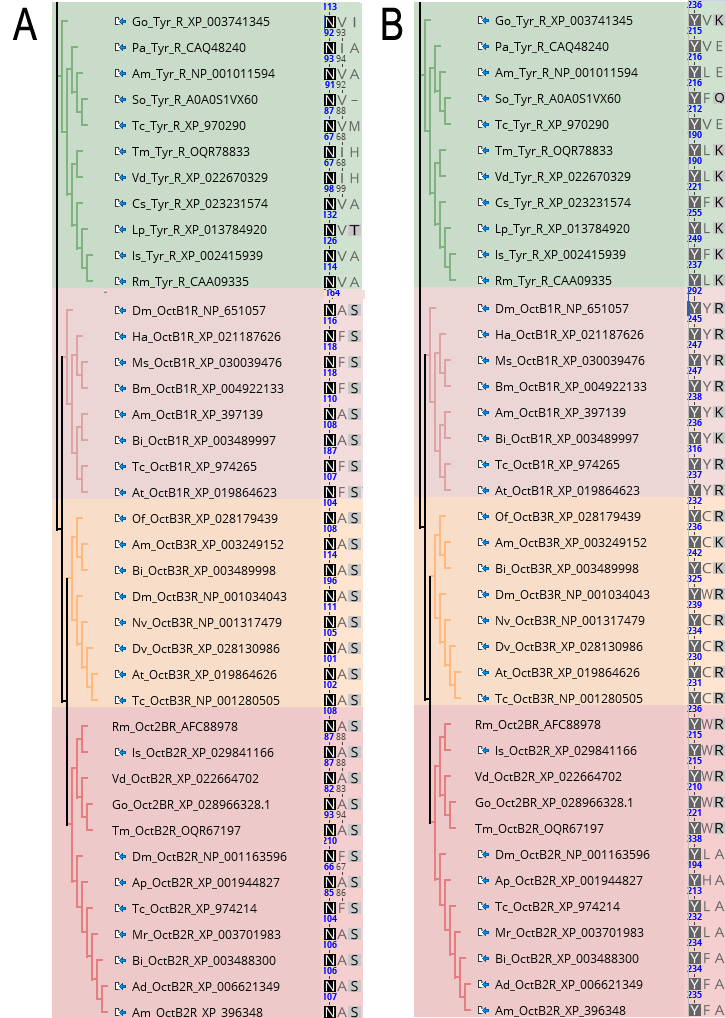
