## Supplementary material for "Resistance to amitraz in the parasitic honey bee mite *Varroa destructor* is associated with mutations in the β−adrenergic-like octopamine receptor": Table S

Table S1. Frequency of the Y215H mutation in the samples collected from several U.S. states.

| Sample | State | Project | n | Last treatment with amitraz | Data collection | % Y215H mutation |
|---|---|---|---|---|---|---|
| NJM-001 | New Jersey | | 24 | 08/2018 | 10/2018 | 50 |
| NJM-002 | New Jersey | | 19 | 09/2018 | 10/2018 | 89 |
| NJM-008 | New Jersey | | 24 | 07/2018 | 10/2018 | 96 |
| EP-2 | New Jersey | | 24 | 04/2018 | 10/2018 | 54 |
| NJ-03 | New Jersey | NHBDS | 24 | | 06/2016 | 0 |
| NJ-08 | New Jersey | NHBDS | 24 | | 09/2016 | 0 |
| NJ-11 | New Jersey | NHBDS | 20 | | 04/2016 | 0 |
| HI-OR-AV01 | Oregon | BIP | 24 | 09/2018 | 10/2018 | 88 |
| HI-OR-AV02 | Oregon | BIP | 23 | 09/2018 | 09/2018 | 96 |
| HI-OR-AL38 | Oregon | BIP | 16 | | 09/2018 | 100 |
| HI-OR-AL51 | Oregon | BIP | 16 | | 09/2018 | 96 |
| HI-OR-1681 | Oregon | BIP | 8 | | 07/2017 | 60 |
| HI-OR-1685 | Oregon | BIP | 16 | | 07/2017 | 31 |

| | | | | | | |
|---|---|---|---|---|---|---|
| OR-24 | Oregon | NHBDS | 13 | | 06/2017 | 0 |
| KP-MI-22 | Michigan | BIP | 17 | 09/2018 | 10/2018 | 94 |
| KP-MI-33 | Michigan | BIP | 17 | 09/2018 | 10/2018 | 90 |
| KP-MI-56 | Michigan | BIP | 24 | | 10/2018 | 100 |
| KP-MI-58 | Michigan | BIP | 23 | | 10/2018 | 96 |
| MI-19 | Michigan | NHBDS | 16 | 09/2017 | 10/2017 | 63 |
| MI-04 | Michigan | NHBDS | 16 | | 10/2016 | 0 |
| MI-09 | Michigan | NHBDS | 24 | | 06/2016 | 10 |
| MA-10 | Massachusetts | NHBDS | 32 | | 2020 | 19 |
| MA-13 | Massachusetts | NHBDS | 15 | | 08/2017 | 33 |
| MA-22 | Massachusetts | NHBDS | 16 | | 08/2016 | 0 |
| MA-21 | Massachusetts | NHBDS | 16 | | 07/2016 | 0 |
| MT-13 | Montana | NHBDS | 28 | | 2020 | 93 |

| | | | | | |
|---|---|---|---|---|---|
| MT-20 | Montana | NHBDS | 16 | 09/2017 | 81 |
| MT-03 | Montana | NHBDS | 16 | 07/2017 | 0 |
| DE-22 | Delaware | NHBDS | 17 | 2020 | 6 |
| DE-13 | Delaware | NHBDS | 14 | 11/2016 | 0 |
| DE-19 | Delaware | NHBDS | 15 | 11/2016 | 0 |
| PA-11 | Pennsylvania | NHBDS | 20 | 2020 | 5 |
| PA-13 | Pennsylvania | NHBDS | 14 | 12/2016 | 0 |
| PA-07 | Pennsylvania | NHBDS | 15 | 11/2016 | 0 |

Table S2. Octopamine and tyramine receptor sequences in arthropod species.

| Species | Protein description | Accession | Length | Abbreviation |
|---|---|---|---|---|
| *Apis mellifera* | Octopamine receptor | NP_001011565 | 587 aa | Am_OctAR |
| *Bombyx mori* | Octopamine receptor | NP_001091748 | 507 aa | Bm_OctAR |
| *Galendromus occidentalis* | Probable G-protein coupled receptor No9 | XP_028966904 | 507 aa | Go_OctAR |
| *Ixodes scapularis* | Probable G-protein coupled receptor No9 | XP_029851527 | 533 aa | Is_OctAR |
| *Rhipicephalus microplus* | Alpha 2 adrenergic-like octopamine receptor | AFC88977 | 480 aa | Rm_OctAR |
| *Tribollium castaneum* | Probable G-protein coupled receptor No9 | NP_001280520 | 586 aa | Tc_OctAR |
| *Tropilaelaps mercedesae* | octopamine receptor 1-like | OQR70152 | 584 aa | Tm_OctAR |
| *Varroa destructor* | Probable G-protein coupled receptor No9 | XP_022669321 | 532 aa | Vd_OctAR |
| *Apis mellifera* | Octopamine receptor beta-1R | XP_397139 | 427 aa | Am_Octβ1R |
| *Aethina tumida* | Octopamine receptor beta-1R-like | XP_019864623 | 447 aa | At_Octβ1R |
| *Bombus impatiens* | Octopamine receptor beta-1R-like | XP_003489997 | 427 aa | Bi_Octβ1R |

| | | | | |
|---|---|---|---|---|
| *Bombyx mori* | Octopamine receptor beta-1R | XP_004922133 | 444 aa | Bm_Octβ1R |
| *Drosophila melanogaster* | Octopamine beta1 receptor, isoform A | NP_651057 | 508 aa | Dm_Octβ1R |
| *Helicoverpa armigera* | Octopamine receptor beta-1R-like | XP_021187626 | 442 aa | Ha_Octβ1R |
| *Manduca sexta* | Octopamine receptor beta-1R-like | XP_030039476 | 444 aa | Ms_Octβ1R |
| *Tribollium castaneum* | Octopamine receptor beta-1R | NP_001280514 | 527 aa | Tc_Octβ1R |
| *Apis dorsata* | Octopamine receptor beta-2R-like isoform X3 | XP_006621349 | 412 aa | Ad_ Octβ2R |
| *Apis mellifera* | Octopamine receptor beta-2R isoform X1 | XP_396348 | 438 aa | Am_ Octβ2R |
| *Acyrthosiphon pisum* | Octopamine receptor beta-2R | XP_001944827 | 443 aa | Ap_ Octβ2R |
| *Bombus impatiens* | Octopamine receptor beta-2R-like isoform X1 | XP_003488300 | 437 aa | Bi_Octβ2R |
| *Drosophila melanogaster* | Octopamine beta2 receptor, isoform F | NP_001163596 | 630 aa | Dm_Octβ2R |
| *Galendromus occidentalis* | Octopamine receptor beta-2R-like | XP_028966328 | 398 aa | Go_Octβ2R |
| *Ixodes scapularis* | Octopamine receptor beta-2R isoform X1 | XP_029841166 | 384 aa | Is_Octβ2R |

| | | | | |
|---|---|---|---|---|
| *Megachile rotundata* | Octopamine receptor beta-2R-like isoform X1 | XP_003701983 | 435 aa | Mr_ Octβ2R |
| *Rhipicephalus microplus* | Beta 2 adrenergic-like octopamine receptor | AFC88978 | 389 aa | Rm_Octβ2R |
| *Tribollium castaneum* | Octopamine receptor beta-2R isoform X2 | XP_974214 | 392 aa | Tc_Octβ2R |
| *Tropilaelaps mercedesae* | Octopamine receptor beta-3R-like | OQR67197 | 439 aa | Tm_Octβ?R |
| *Varroa destructor* | Octopamine receptor beta-2R-like isoform X6 | XP_022664702 | 366 aa | Vd_Octβ2R |
| *Aethina tumida* | Octopamine receptor beta-3R-like | XP_019864626 | 397 aa | At_Octβ3R |
| *Apis mellifera* | Octopamine receptor beta-3R isoform X1 | XP_003249152 | 413 aa | Am_Octβ3R |
| *Bombus impatiens* | Octopamine receptor beta-3R-like isoform X1 | XP_003489998 | 419 aa | Bi_Octβ3R |
| *Drosophila melanogaster* | Octopamine beta3 receptor, isoform J | NP_001034043 | 445 aa | Dm_Octβ3R |
| *Diabrotica virgifera virgifera* | Octopamine receptor beta-3R-like | XP_028130986 | 435 aa | Dv_Octβ3R |
| *Nicrophorus vespilloides* | Octopamine receptor beta-3R-like | NP_001317479 | 446 aa | Nv_Octβ3R |
| *Ostrinia furnacalis* | Octopamine receptor beta-3R-like | XP_028179439 | 445 aa | Of_Octβ3R |

| | | | | |
|---|---|---|---|---|
| *Tribollium castaneum* | Octopamine receptor beta-3R-like | NP_001280505 | 398 aa | Tc_Octβ3R |
| *Apis mellifera* | Tyramine receptor | NP_001011594 | 399 aa | Am_TyrR |
| *Centruroides sculpturatus* | Probable G-protein coupled receptor No18 | XP_023231574 | 405 aa | Cs_TyrR |
| *Galendromus occidentalis* | Tyramine receptor 1-like | XP_003741345 | 425 aa | Go_TyrR |
| *Ixodes scapularis* | Tyramine receptor 1 | XP_002415939 | 400 aa | Is_TyrR |
| *Limulus polyphemus* | Probable G-protein coupled receptor No18 | XP_013784920 | 439 aa | Lp_TyrR |
| *Periplaneta americana* | Tyramine receptor 1 | CAQ48240 | 441 aa | Pa_TyrR |
| *Rhipicephalus microplus* | G-protein coupled receptor | CAA09335 | 419 aa | Rm_TyrR |
| *Sitophilus oryzae* | Octopamine receptor | XP_030755585 | 455 aa | So_TyrR |
| *Tribollium castaneum* | Tyramine/octopamine receptor | NP_001164311 | 453 aa | Tc_TyrR |
| *Tropilaelaps mercedesae* | G-protein coupled receptor-like | OQR78833 | 369 aa | Tm_TyrR |
| *Varroa destructor* | Octopamine receptor-like isoform X1 | XP_022670329 | 369 aa | Vd_TyrR |

Table S3. Primers used for amplification and sequencing of *Vd_octα₂r*, *Vd_octβ₂r* and *Vd_tar1* genes.

| **Gene** | | **Name** | **Sequence 5'→ 3'** | **Localization** |
|---|---|---|---|---|
| *Vd_octα₂r* | Forward primers | Vd_OctAR_5UTR | CAACATCGGTCGTTTCACTG | 5'UTR |
| | | Vd_OctAR_344F | GCCACTTCTGCAAGATCTGG | Exon 6 |
| | | Vd_OctAR_881F | TGGCCAAGAACAAAGGTGGA | Exon 7 |
| | Reverse primers | Vd_OctAR_938R | CGCAGGGTCATTTGTTGCAT | Exon 7 |
| | | Vd_OctAR_3UTR | GGACTAAACACTGCGGGGTA | 3'UTR |
| *Vd_octβ₂r* | Forward primers | Vd_OctBR_5UTR1 | CACGAACGACAAAACCGGTC | 5'UTR |
| | | Vd_OctBR_5UTR3 | GCGTCGGTTGAAATCGGAAG | 5'UTR |
| | | Vd_OctBR_246F | CGCGATGACATTCAATGCGT | Exon 1 |
| | | Vd_OctBR_476F | GCTCATATCCTTCGTGCCCA | Exon 1 |
| | | Vd_OctBR_iF | CGTTGATGTCTGTTTGCTGTTTG | Intron |
| | Reverse primers | Vd_OctBR_563R | GACCATCCGAACGAGTGTGT | Exon 1 |

| | | | | |
|---|---|---|---|---|
| | | Vd_OctBR_437iR | AGCTACGCATGCTGATCGAT | Intron |
| | | Vd_OctBR_1103R | GATTTGTGAAAGTAAAGTGTGACG | Exon 2 |
| | | Vd_OctBR_3UTR | TAATCACGCCCACGGATACG | 3’UTR |
| *Vd_tar1* | Forward primers | Vd_TAR1_5F | GGTCTATCTCATTTTCTTTTCACCG | 5’UTR |
| | | Vd_TAR1_E3F | CCATAAACGCACTTTACGGCGGGTG | Exon 3 |
| | | Vd_TAR1_i3F | GAGGAGGCTAGTCAGGCCAAGATGGCC | Intron 3 |
| | Reverse primers | Vd_TAR1_E3R | CTTTCTCGGAGCCGTTTTCGCGTTG | Exon 3 |
| | | Vd_TAR1_i3R | GGCCATCTTGGCCTGACTAGCCTCCTC | Intron 3 |
| | | Vd_TAR1_E4R | GCCACCGAATCTTCACGTTCAGCG | Exon 4 |
| | | Vd_TAR1_3R | GAGATTAGGGATGCATCAGACGAACGC | Exon 4-3’UTR |
